# Nanopore sequencing reveals endogenous NMD-targeted isoforms in human cells

**DOI:** 10.1101/2021.04.30.442116

**Authors:** Evangelos D. Karousis, Foivos Gypas, Mihaela Zavolan, Oliver Mühlemann

**Affiliations:** Department of Chemistry, Biochemistry and Pharmaceutical sciences, University of Bern, Freiestrasse 3, Bern, 3012,Switzerland; Friedrich Miescher Institute for Biomedical Research, Maulbeerstrasse 66, Basel 4058, Switzerland; Biozentrum, University of Basel and Swiss Institute of Bioinformatics, Klingelbergstrasse 50-70, 4056 Basel, Switzerland

**Keywords:** Nonsense-mediated mRNA decay, NMD, Long-read sequencing, Nanopore sequencing, mRNA degradation, cDNA sequencing, transcriptomics, mRNA isoforms

## Abstract

**Background:** Nonsense-mediated mRNA decay (NMD) is a eukaryotic, translation-dependent degradation pathway that targets mRNAs with premature termination codons and also regulates the expression of some mRNAs that encode full-length proteins. Although many genes express NMD-sensitive transcripts, identifying them based on short-read sequencing data remains a challenge.

**Results:** To identify and analyze endogenous targets of NMD, we applied cDNA Nanopore sequencing and short-read sequencing to human cells with varying expression levels of NMD factors. Our approach detects full-length NMD substrates that are highly unstable and increase in levels or even only appear when NMD is inhibited. Among the many new NMD-targeted isoforms that our analysis identified, most derive from alternative exon usage. The isoform-aware analysis revealed many genes with significant changes in splicing but no significant changes in overall expression levels upon NMD knockdown. NMD-sensitive mRNAs have more exons in the 3΄UTR and, for those mRNAs with a termination codon in the last exon, the length of the 3΄UTR *per se* does not correlate with NMD sensitivity. Analysis of splicing signals reveals isoforms where NMD has been co-opted in the regulation of gene expression, though the main function of NMD seems to be ridding the transcriptome of isoforms resulting from spurious splicing events.

**Conclusions:** Long-read sequencing enabled the identification of many novel NMD-sensitive mRNAs and revealed both known and unexpected features concerning their biogenesis and their biological role. Our data provide a highly valuable resource of human NMD transcript targets for future genomic and transcriptomic applications.

## Background

Ribonucleolytic activities are essential to dispose cells of defective RNAs, protect host cells from infections with RNA viruses, and regulate gene expression (1,2). Nonsense-mediated mRNA decay (NMD) is one of the degradation pathways that in eukaryotes is involved in all these functions, on a broad range of RNA substrates (3,4). NMD was discovered as a mechanism that rids eukaryotic cells of mRNAs with premature termination codons (PTCs) arising from mutations or errors in splicing or transcription. However, subsequent application of transcriptome-wide approaches has revealed that NMD also targets many mRNAs that encode full-length proteins, to regulate their overall expression level. It is not a surprise that by changing the levels of endogenous mRNAs, NMD affects various biological processes which are dependent on the targeted mRNAs (5).

Although many protein factors that recognize and degrade these mRNA substrates have been identified, how the recognition of substrates and NMD activation are accomplished remains unclear. In human cells, after the completion of at least one translation cycle of an NMD-sensitive mRNA (3,6,7), the RNA helicase UPF1 that is bound or recruited on the targeted mRNP is phosphorylated by the phosphatidylinositol-kinase related kinase (PIKK) SMG1 (8). Phosphorylated epitopes of UPF1 form a platform that recruits the endonuclease SMG6 and the adaptor proteins SMG5 and SMG7. SMG6 directly cleaves the RNA near the termination codon (9–11), whereas the SMG5-SMG7 heterodimer recruits general deadenylation and decapping factors that catalyze the degradation of the mRNA (11–14). The NMD activity on individual substrates is modulated by additional factors, and many different models have been put forward to address the mechanistic details of NMD (reviewed in (3,4,15–17)).

While it has been established that NMD has a vital function in ridding transcriptomes of aberrant mRNAs with PTCs, cases in which NMD serves to rapidly switch off the expression of specific proteins have also emerged (18). Alternative splicing coupled to NMD (AS-NMD) is a known mechanism that regulates the concentration of specific mRNA isoforms (19). Examples of this mechanism have been reported for all 11 human genes that encode the arginine-serine-rich (SR) proteins that regulate splicing. The transcripts encoding these proteins have PTC-introducing exons whose splicing depends on the expression level of the corresponding protein, thereby building an autoregulatory feedback loop (19,20). By shifting pre-mRNA splicing towards unproductive transcripts that are degraded by NMD, the expression levels of these abundant proteins decreases (2). The regulatory potential of NMD is illustrated by its targeting of transcripts encoding RNA-binding proteins that modulate the splicing of their own transcripts, creating autoregulatory feedback loops (21). In cancer cells, perturbations in such feedback loops may lead to the expression of neoantigens that contribute to the development of the disease (22,23).

Through its quality control function in degrading aberrant endogenous mRNAs, NMD is also implicated in the development of various diseases. NMD inhibition has been reported in cancer, where it leads to the stabilization of transcripts that are important for tumorigenesis, such as KLF6 (Kruppel-like factor 6) in hepatocellular carcinoma and MALAT1 (metastasis-associated lung adenocarcinoma transcript 1) in gastric cancer (24,25). A synergistic effect between splicing and NMD has been observed for the alternatively spliced isoform β of p53, a gene that is highly relevant to cancer progression (26).

These findings demonstrate that accurate identification of NMD-targeted RNAs is crucial to better understand how transcriptomes are remodeled in various diseases, where transcripts that in normal cells are rapidly degraded and thus have very low expression levels become more stable and alter cellular functions. Comprehensive catalogs of NMD-sensitive transcripts serve as a basis for identifying features that contribute to the recruitment of NMD factors and to their activity. They also enable the identification of regulatory circuits that operate on specific transcripts in different conditions. We and others have performed short-read sequencing from cells with decreased NMD activity to expose endogenous NMD targets (reviewed in (3,15,27)). These approaches have revealed several features of NMD substrates: exons in the 3΄UTR, upstream open reading frames (uORFs), and unusually long 3΄UTR have all been associated with the sensitivity to NMD. Additionally, evidence from cancer cells indicates that exons longer than 400 nt, specific motifs for RNA-binding proteins, and the mRNA half-life may also contribute to the NMD sensitivity of physiological or aberrant mRNAs (28). However, quantifying the abundance of the NMD isoforms with typically low expression from short-read data cannot be done accurately (29) and as a consequence, the catalog of NMD substrates and of their NMD-stimulating features remains incomplete.

A well-known caveat of short-read sequencing is that natural nucleic acid polymers that vary widely in length need to be reassembled from the short reads and quantified. This task is challenging when alternative splicing and polyadenylation lead to the expression of transcript isoforms that differ over only a small proportion of their length (30–32). Long-read sequencing, which currently yields reads that exceed the length of 10 kb, can overcome this caveat to reveal full-length transcript isoforms (33–35). Several studies that have applied either total or targeted long-read sequencing revealed that even the most comprehensive annotations still miss a vast amount of information concerning expressed transcripts (36,37). The current limitation of long-read sequencing, however, is its moderate sequencing depth compared to short-read sequencing, which impedes accurate quantification of less abundant mRNAs.

Previously, we applied short-read RNA-seq to knockdown (KD) and rescue conditions of three NMD factors (UPF1, SMG6 and SMG7) to detect genes with NMD-sensitive isoforms (38). To increase the resolution, depth and accuracy of NMD-sensitive isoform identification, we here combined long-read Nanopore cDNA sequencing with short-read sequencing. We used the long reads to create a curated reference transcriptome, while the short reads were used to estimate the abundance of individual isoforms and identify those that respond to fluctuations in the levels of NMD factors. Splicing analysis of the NMD-transcriptome showed that many NMD targets derive from alternative exon events, but other types of splicing isoforms that would be difficult to resolve solely based on short reads are also detected. Our data highlights the central role of exon junctions in the 3΄UTR as an NMD-triggering feature. Interestingly and in contrast to previous reports, for mRNAs with a termination codon in the last exon, the length of the 3΄UTR is not correlated with NMD sensitivity, highlighting again the importance of analyzing full-length transcript isoforms. We also revealed that NMD targets canonically and non-canonically spliced mRNAs, indicating that NMD serves as a regulatory mechanism but also as a mechanism to rid the transcriptome of aberrantly spliced transcripts.

## Results

### NMD Inactivation and Nanopore cDNA sequencing

In a previous study, we set out to identify NMD targets by Illumina sequencing. We provided a list of genes whose expression is sensitive to NMD, but the data did not allow us to unambiguously identify the NMD targets at isoform level (38). To resolve this issue and thereby improve the coverage and accuracy of NMD isoform identification, we here employed long-read sequencing. We depleted UPF1, SMG6, or SMG7 proteins individually in HeLa cells using shRNA-mediated KD and compared their isoform expression with that of cells subjected to control KD (CTR) using an shRNA with a scrambled sequence. In a second set of experiments, we knocked down SMG6 and SMG7 alongside another control experiment. The KD efficiency was validated by western blotting, as shown in Fig. 1A.

**Figure 1.**
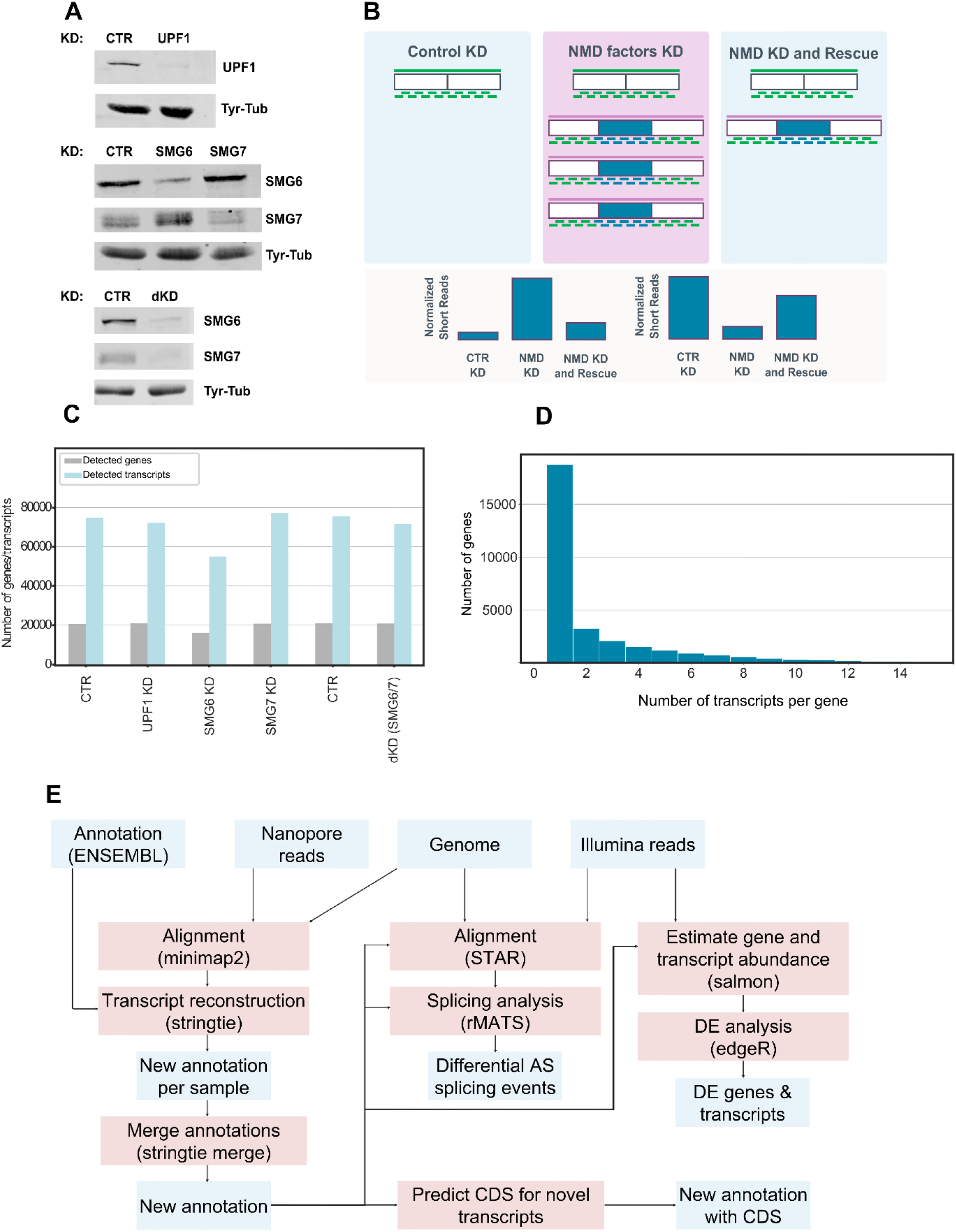
**(A)** Western blot analysis of HeLa cell lysates corresponding to 2×10^5^ cell equivalents of cells transiently transfected with the indicated knockdown constructs. Membrane sections were incubated with antibodies against UPF1, SMG6, SMG7, and Tyr-Tubulin, the latter serving as a loading control. **(B)** Upper part: Schematic representation of how long and short-read sequencing are combined to identify endogenous NMD-sensitive mRNA isoforms in human cells. Boxes denote exons (NMD-inducing exons in blue,), green lines denote long and short sequencing reads, long purple lines denote long reads that correspond to NMD-sensitive isoforms, short blue lines denote short reads that map to exons of NMD-sensitive isoforms. Lower part: Representation of the short-reads expression level patterns of NMD-sensitive exons. NMD-sensitive isoforms can occur by exon inclusion or exon exclusion and the patterns of changes of the expression levels to opposite directions are taken into consideration. **(C)** Bar plot of the number of genes and transcripts detected by long-read cDNA sequencing in different experimental conditions. **(D)** Histogram depicting the number of isoforms that were detected per gene cumulated over all the conditions. **(E)** Schematic illustration of the bioinformatics pipeline for analyzing NMD-sensitive mRNA isoforms using long and short-read data. The components of the pipeline shaded in light blue describe input/output files and the boxes in red represent the computational tools that were applied.

Figure 1B describes the concept of our approach schematically. Under normal conditions, when NMD is functional, RNAs targeted by NMD have low or even undetectable expression. When the NMD activity is reduced, NMD-sensitive transcripts accumulate and can be detected by long-read sequencing. We, therefore, used the long reads to create a curated transcriptome that contains NMD-sensitive isoforms and serves as a reference for mapping short sequencing reads and quantifying alternative splicing isoforms that are absent from other annotations (Figure 1B). This approach allows the comparative analysis of RNAs that are targeted by NMD at an isoform-specific level.

We extracted total RNA from cells, isolated polyA+ mRNAs, prepared cDNA libraries according to the Nanopore protocol and carried out direct cDNA sequencing. Nanopore sequencing was performed on a GridION using R9.4 flowcells, and the MinKNOW instrument software was used to record the Nanopore current. Basecalling was performed using GUPPY version 1.4.3-1 from Oxford Nanopore Technologies.

The 0.8 to 2.4 million long reads that were obtained from the different samples (Sup. Fig.1A) were aligned to the human reference genome using minimap2 (39), with the ENSEMBL reference annotation (40). 17’000 to 22’000 genes had evidence of expression across samples (Sup. Fig.1). 55’000-77’000 different isoforms were detected in the samples (Fig.1C) and all samples showed a similar size distribution of the read length (Sup. Fig. 1B). Overall, an average of 3.4 transcripts per gene was detected (Fig.1D). Given that most isoforms were present in very few copies in a sample, we decided to combine long sequencing data with Illumina sequencing data to quantify splicing events that give rise to NMD-sensitive mRNAs.

The short reads originated from a previous sequencing study in our lab that was performed under the same experimental conditions after knocking down the same NMD factors in triplicates. In this work, we analyzed by Illumina sequencing three biological replicates from HeLa cells under ten different treatment conditions: control, UPF1, SMG6 or SMG7 single KD, single KD and rescue of each of these factors, or double KD (dKD) of SMG6 and SMG7 accompanied by rescues with SMG6 or SMG7 individually (38).

The bioinformatics pipeline for the analysis of NMD-sensitive mRNAs is shown in Fig. 1E. We used *StringTie2* (41) to create an augmented reference transcriptome that contains not only the ENSEMBL transcriptome but also the novel isoforms that were identified by long-read sequencing. Short reads were aligned with *STAR* (Spliced Transcripts Alignment to a Reference) to this reference transcriptome to find covered splice junctions (42). The splicing analysis was carried out with Replicate Multivariate Analysis of Transcript Splicing *(rMATs)* (43). We used *Salmon* (44) to calculate expression levels of genes and transcripts by edgeR (45) for the differential expression analysis.

### NMD targets specific RNA isoforms

We first assessed to what extent NMD inhibition alters the expression of alternative transcript isoforms. We applied rMATS, a statistical model that allows detection of differential alternative splicing events from two groups of RNA-Seq samples. rMATS quantifies the usage of various types of isoforms in the “percent spliced in” (PSI) variable, indicating the proportion of transcripts that provide evidence for the inclusion of a specific alternatively spliced region among all pre-mRNAs that cover that region and taking into consideration the variability among the replicates (43).

To further increase the chance of identifying functionally relevant events that react to NMD modulation, we selected those cases that (i) showed at least 10% change upon KD of NMD factors and (ii) changed in the opposite direction when the corresponding NMD factor was overexpressed (rescue experiment). The types of events that were analyzed are depicted schematically in Fig. 2A. As seen in Fig. 2B, the most common event in our data set of NMD-sensitive transcripts was the introduction of alternative exons, which refers to NMD-sensitive transcripts arising by an exon being either included or excluded compared to the NMD-insensitive isoform. Out of 4’137 skipping exon events that give rise to NMD-sensitive isoforms, 2’490 represent exon inclusion and 1’647 exon exclusion events. Interestingly, we observed several cases of genes with more than one splicing events leading to NMD, because the number of the splicing events is higher than the number of the affected genes (Fig.2B). We also identified events involving mutually exclusive exons, alternative 5΄ and 3΄ splice sites, as well as retained introns, but these were less frequently compared to alternative exon events (Sup. Fig. 2A-E).

**Figure 2.**
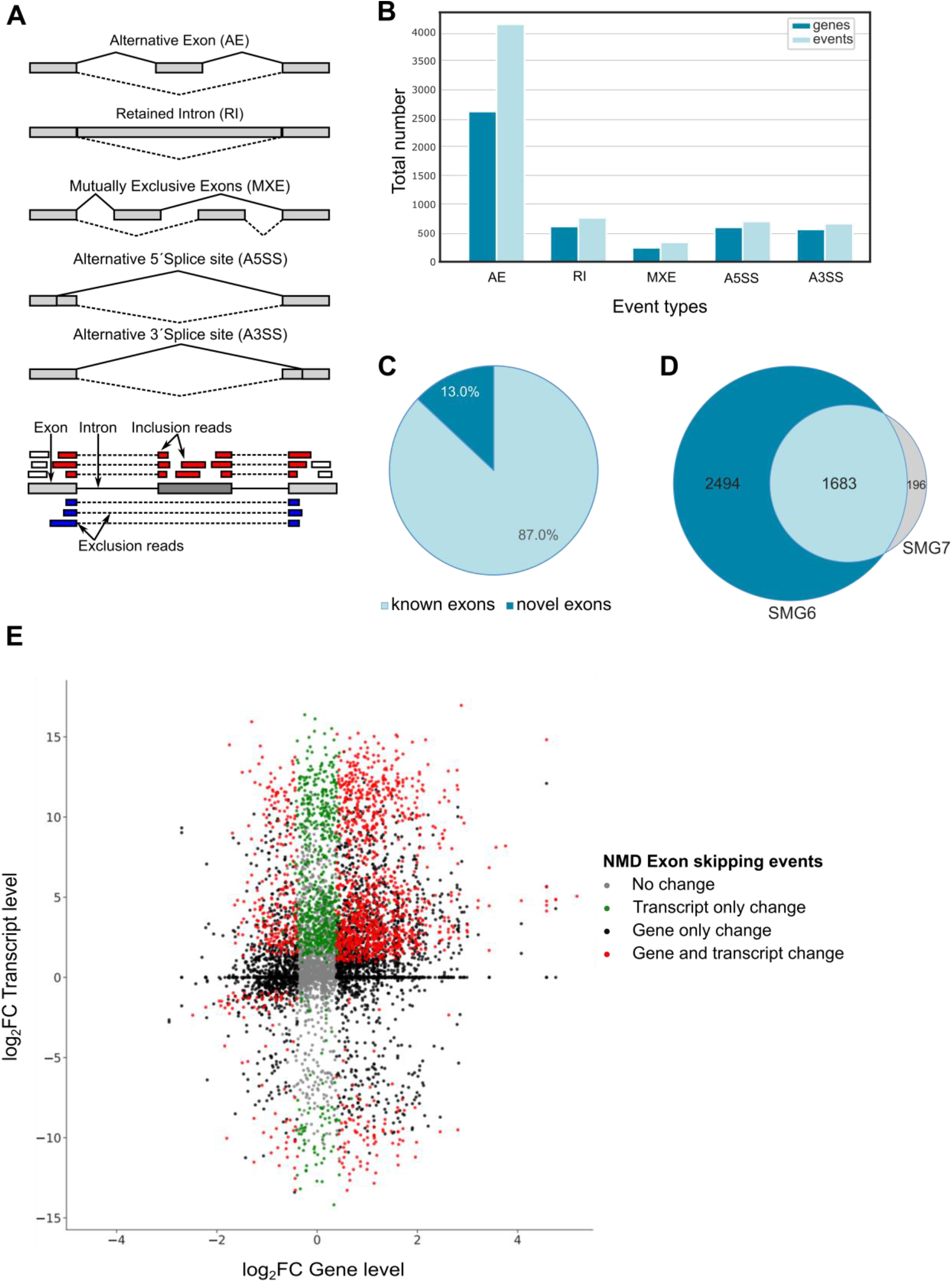
**(A)** Upper part: Schematic representation of the types of splicing events analyzed in this study. Lower part: Schematic representation of short-read assignment to the reference transcriptome for skipping exon events. **(B)** Bar plot depicting the number of different splicing events that follow the expected pattern of expression of NMD targets. **(C)** Pie chart representing the percentage of novel (identified by long-read cDNA sequencing) and previously known exons (contained in the previous annotation) that are represented in the augmented annotation. **(D)** Pie chart showing the sets of exons found to respond to SMG6 or SMG7 KD. **(E)** Scatter plot depicting the log2 fold change (log2FC) of NMD-sensitive transcripts (y-axis) versus the log2FC between dKD and CTR of their corresponding genes (x-axis) showing alternative exon events (Ν= 8’056) based on SMG6/SMG7 double KD, CTR KD and SMG6 Rescue conditions.

To assess whether the usage of long reads facilitated the identification of novel NMD-sensitive isoforms, we calculated the percentage of exons that are included in our augmented human transcriptome but absent in the currently annotation. As shown in Fig. 2C, among the events with significant splicing changes, approximately 13% are only present in the sequences originating from the Nanopore sequencing reads. Therefore, by augmenting the reference transcriptome with long read-supported isoforms, we identified numerous previously unknown splicing events.

To further validate our approach, we compared the set of events that responded to either SMG6 or SMG7 rescue of cells with dKD of both of these factors. As shown in Fig. 2D, we observed a large overlap between the two sets of transcript isoforms, validating our approach and further demonstrating the redundancy of the two branches of NMD that we observed and reported before (38).

Next, we addressed the power of our approach in identifying NMD-sensitive isoforms beyond those found before with short-read sequencing. Short-read-based quantification of isoform expression depends on an accurate reference transcriptome, which is difficult to reconstruct from the short reads alone. Therefore, prior studies relied on gene-level quantification to detect cases that responded to perturbations in NMD factors. To illustrate the utility of the long-read-based approach, we asked whether there are changes in the expression of transcript isoforms that are not detected at the gene level. For this analysis, we focused on the most abundant type of NMD-inducing events, skipping exons, and in particular those that show a higher exon inclusion level upon dKD compared to control and whose level is again reduced by the rescue with SMG6. As shown in Fig. 2E, while the KD of NMD factors brings about broad changes in gene expression, about 30% of the genes deemed as differentially expressed in dKD vs. CTR (1’488 of 5’795) have isoforms whose levels are also significantly changed by NMD perturbation. In contrast, 686 of 8’056 of the genes for which no significant change is detected at the gene level, have an isoform that changes significantly in dKD relative to CTR, likely because the NMD-sensitive isoform forms a small proportion of all the transcripts emanating from that gene. These cases are missed by pipelines that select NMD targets on the basis of differential expression at the gene level.

### Validation of new NMD-sensitive isoforms

The apoptosis-modulating Bcl-2-associated athanogene-1 (BAG-1) has three main protein isoforms, deriving from alternative translation initiation on a single mRNA and showing different localization patterns (46). It has been reported that BAG-1 mRNAs are stabilized upon NMD depletion because of the presence of a uORF that renders these mRNAs NMD-sensitive (47). Long-read sequencing revealed an alternative isoform that includes an alternative exon and is stabilized upon NMD inhibition (Fig. 3A). To verify the expression of this isoform in the cells, we designed PCR assays that target specifically either the protein-coding isoform or the NMD-sensitive isoform. As shown in Fig. 3A, the protein-coding isoform is clearly visible in both conditions (CTR and dKD), whereas the NMD-sensitive isoform that harbors an NMD-causing exon showed a clear stabilization upon NMD inhibition.

**Figure 3.**
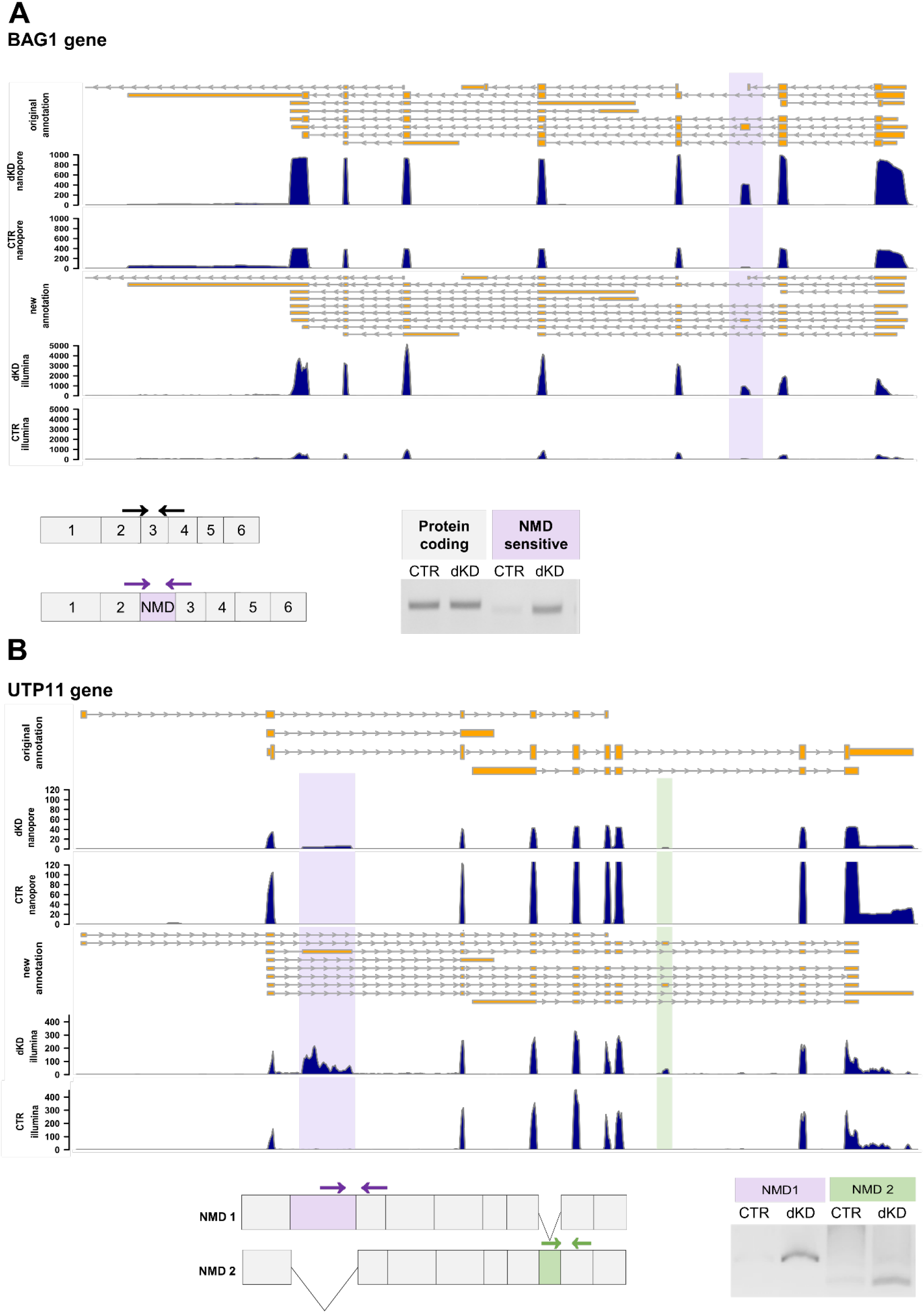
**(A and B)** Genome browser tracks at the top of each panel show the intron (grey lines) / exon (yellow boxes) annotations of the different transcripts of the BAG1 (A) and UTP11 (B) genes according to ENSEMBL (original annotation) and according to the new transcriptome expanded by the long-read data (new annotation). Arrows denote the direction of transcription. The coverage of long (Nanopore) and short reads (Illumina) of the control (CTR) and SMG6/SMG7 double knockdown (dKD) samples is shown in dark blue. In the scheme underneath the genome browser tracks, constitutive exons are depicted by grey boxes, those that induce NMD by purple and light green boxes. The relative positions of primers used for splice-isoform-specific RT-PCR are shown by arrows. Agarose gels show the products of PCR reactions using the designated primers.

UTP11 is an example that illustrates the advantages of our approach to study NMD-sensitive mRNA isoforms using long-read sequencing. The UPT11 gene encodes the human homolog of U3 snoRNA-associated protein 11 that was identified in yeast and is implicated in the nucleolar processing of pre-18S ribosomal RNA (48). Our data revealed a clear enrichment of two so-far unannotated exons. These are included in two different transcripts (Fig. 3B), both of which were detected under decreased cellular NMD activity. To confirm the expression of these isoforms, we designed primers that are specific for amplifying each of these isoforms (NMD isoforms 1 and 2, respectively) by RT-PCR. As shown in Fig 3B, a single band corresponding to each isoform was amplified under dKD conditions, which verifies the inclusion of the poised exons in mRNA isoforms that are specifically targeted by NMD. That no additional PCR products were detected in the assay that is specific for NMD isoform 1 agrees with the long-read sequencing data, which indicates that the two poised exons are included in two different isoforms. This analysis leads to the conclusion that the UTP11 gene encodes two distinct NMD-sensitive mRNA isoforms. Importantly, the attribution of poised exons to specific isoforms would not have been possible without the annotation derived from the long reads.

### Splicing events in the 3΄UTR trigger NMD independently of 3΄UTR length

The features of NMD-sensitive mRNAs can guide the identification of molecular mechanisms of substrate recognition and can clarify the biological role of the NMD pathway in human cells. As our previous efforts to perform a transcript-specific analysis using only short reads were only partially successful, we address this caveat here, using the long read-based augmented transcriptome.

As in general, multiple isoforms provide evidence for both inclusion and skipping of NMD-sensitive exons, the numbers varying widely between genes, we first constructed a non-redundant set of transcript pairs. First, we identified all transcripts related to significant alternative exon events that were enriched upon NMD inhibition and filtered out those without annotated coding regions. Next, we identified NMD-sensitive and NMD-insensitive transcripts providing evidence for a particular splicing event (Supplementary table 1). When multiple isoforms were detected, we selected only the one with the highest expression level (see methods). NMD is a translation-dependent process, and most of the features that render mRNAs sensitive to NMD depend on the relative position of the termination codon within the transcript (4,49). To investigate potential differences in the molecular architecture of NMD-sensitive versus NMD-insensitive mRNAs, we extrapolated the position of the termination codon of NMD-sensitive mRNAs using as a reference the initiation codon of the principal protein isoform of the gene (50). To this end, we used the APPRIS database to annotate the principal protein-coding isoforms, which takes into consideration structural, functional and conservation data (51). Then, by using the start codon of the principal isoform, we determined the termination codon to define the open reading frames of both NMD-sensitive and NMD-insensitive transcripts, as well as the 5΄ and the 3΄ UTRs.

The best-known NMD-inducing feature is the presence of an exon-exon junction more than 55 nucleotides downstream of the termination codon (28,52–55). To assess whether this feature characterizes our NMD-sensitive mRNAs, we compared the number of exon junctions downstream of the termination codon between NMD-sensitive and NMD-insensitive mRNAs, as described above. As shown in Fig. 4A, there is a considerable enrichment of transcripts with an exon junction in the 3΄UTR among the NMD-sensitive mRNAs. More specifically, 80% of the NMD-sensitive transcripts contain an exon junction in the 3΄UTR, whereas only 31% of the NMD-insensitive mRNAs have this feature. This result adds additional confidence in the set of NMD-sensitive transcripts that we identified here. Nevertheless, there are also some isoforms that are sensitive to NMD and that do not contain exon junctions in the 3΄UTR (389 out of 1’911).

**Figure 4.**
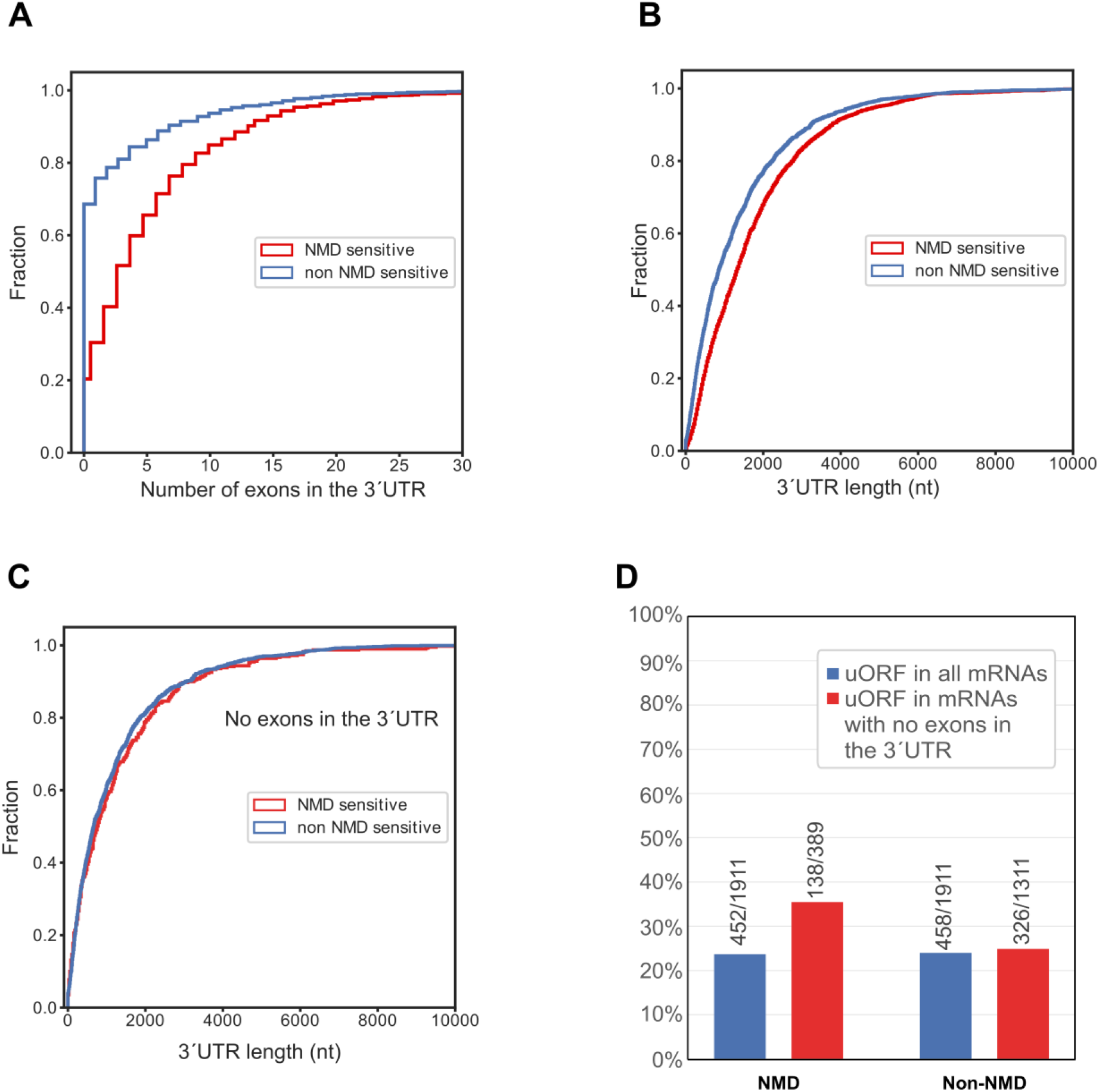
**(A)** Cumulative distribution function (CDF) of the fraction of NMD-sensitive (red) and NMD-insensitive transcript isoforms (blue) with specified numbers of exons in the 3΄UTR. The statistical significance of the difference between the two distributions was determined by the K-S test, *p*=7×10^−202^, N=1’911 **(B)** CDF of the fraction of NMD-sensitive and NMD-insensitive transcript isoforms with specified 3’UTR length. Statistical significance was determined by K-S test, *p*=4×10^−91^, N=1’911 **(C)** CDF of the fraction of NMD-sensitive and NMD-insensitive transcript isoforms with a specified length of the 3΄UTR, restricted to isoforms with an APPRIS-designated termination codon in the last exon. Statistical significance was determined by K-S test, *p*=0.36, N=389 for NMD sensitive and N=1’311 for NMD-insensitive transcripts. **(D)** Relative percentage of experimentally validated uORFs in NMD-sensitive and NMD-insensitive mRNA isoforms computed separately for all isoforms (blue) and for those with a termination codon in the last exon (red). The numbers of transcripts in the different categories are indicated on the plot.

As this is the first long-read analysis from cells with reduced NMD activity, we decided to address the longstanding question of whether NMD also targets endogenous mRNAs with long 3΄UTRs, independently of exon junctions. Comparing NMD-sensitive with NMD-insensitive isoforms, we found that indeed NMD-sensitive mRNAs tend to have, on average, a longer 3΄UTR (Fig. 4B). However, this effect is due to longer 3΄UTRs also containing more exon junctions. When we removed from the list of NMD-sensitive transcripts those with more than one exon in the 3’UTR and retained only those harboring the stop codon in the last exon, we found no difference in the average 3’UTR length between NMD-sensitive and insensitive mRNAs (Fig. 4C). This indicates that the length of the 3΄UTR as an NMD-activating feature in human cells should be revisited.

### The presence of uORFs predicts a subset of NMD-sensitive mRNAs

The translation of upstream open reading frames (uORFs) has also been reported to lead to NMD (3,52,56). To assess this, we identified computationally ORFs starting with an AUG and encoding at least five amino acids (15 nt) within the 5΄UTR and upstream of the main APPRIS-designated ORFs and included those in our analysis, for which a recently published ribosome profiling data set provided experimental evidence for their translation (57). While no difference in the fraction of mRNAs with uORFs can be detected between all NMD-sensitive and insensitive mRNAs, an enrichment among NMD-sensitive mRNAs becomes evident when only mRNAs are considered that do not have exon-exon junctions in their 3’UTR (35.5% versus 24.9%; Fig. 4D). This result indicates that uORFs and exon-exon junctions downstream of the stop codon are orthogonal features conferring NMD-sensitivity. That is, uORFs account for NMD-sensitivity of mRNAs that lack exon-exon junctions in the 3’UTR.

### NMD targets products of canonical and non-canonical splicing

Splicing is a noisy process (58), which often leads to spurious, PTC-containing isoforms. While it is clear that NMD can act as a gene expression regulation mechanism by targeting specific alternatively spliced transcript isoforms or mRNAs that arise from nonsense mutations, to date, it is not clear whether NMD contributes significantly to the degradation of aberrantly spliced mRNAs due to splicing errors. Reasoning that spuriously used splice sites are not under evolutionary selection, we sought to assess to what extent NMD is co-opted in the regulation of gene expression as opposed to clearing out aberrant transcripts by quantifying the strength and conservation of splicing signals among all NMD-sensitive, poised exons. We quantified the strength of splice signals with the MaxENT method (59) and the degree of evolutionary conservation by the average PhyloP 30-way vertebrate conservation score (60). As control exons, we used an equal number of randomly selected exons that we did not find to respond to NMD perturbations. This combined analysis showed differences, mostly in the conservation rate but also in the splicing consensus score of the 3΄ (Fig. 5A) and 5΄ (Fig. 5B) splice sites of NMD-triggering exons and the control group. NMD-sensitive exons have on average weaker and less conserved splice sites compared to control exons. Interestingly however, a small subpopulation of NMD-sensitive exons with stronger splice sites and a higher degree of conservation compared to control exons is also distinguishable. The known poised exons of SR proteins are in this latter category (Fig. 5), indicating that this category contains other exons with important regulatory functions. These results further imply that NMD exerts a dual role in shaping the transcriptome, the primary function being the clearance of transcripts resulting from aberrant splicing involving weak and poorly conserved splice sites. In addition, NMD also plays a role in the regulation of gene expression through the degradation of transcripts with highly conserved alternative exons.

**Figure 5.**
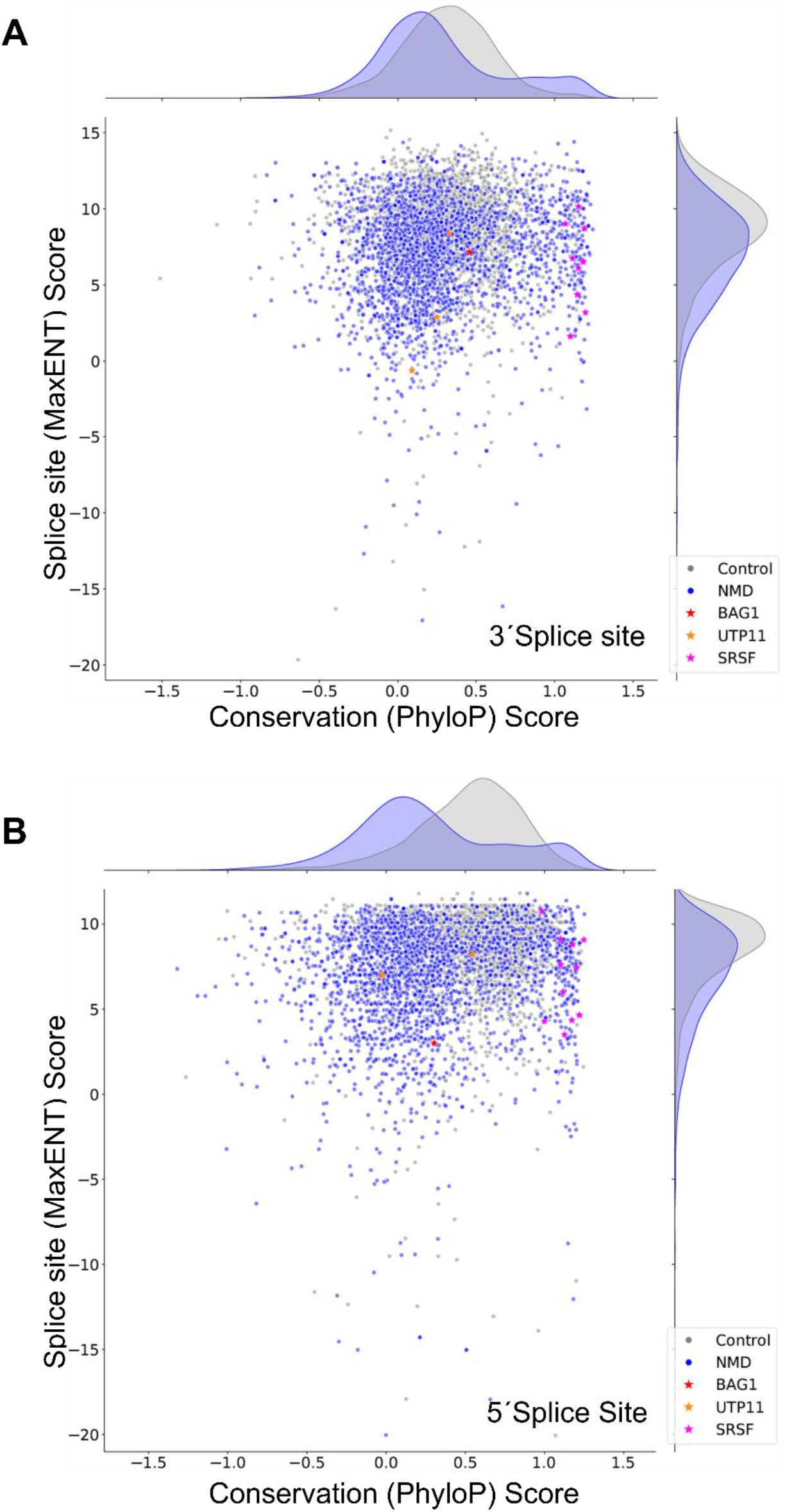
Combined scatter plots of MaxENT scores (denoting splice site strength) and PhyloP 30-way mean conservation scores for the 3΄ (Α) and 5΄ (B) splice sites of an equal number of poised, NMD-triggering exons (blue - NMD) and constitutive exons (grey - Control) (N=2’405). NMD-inducing exons of the genes BAG1, UTP11 and SR-encoding proteins are indicated.

## Discussion

In this work, we applied Nanopore long-read sequencing in combination with Illumina short-read sequencing to identify endogenous human mRNAs that are targeted by NMD. By depleting NMD factors, we stabilized mRNA isoforms that in normal conditions are rapidly degraded by NMD and, therefore, difficult to detect in cells with normal NMD activity. This isoform-specific analysis showed that NMD-sensitive mRNAs derive mostly from the inclusion of alternative exons that have either canonical or non-canonical splice sites, indicating that NMD has evolved to serve two roles. As a quality control system, it can recognize and eliminate aberrantly processed transcripts, and as a posttranscriptional gene regulation mechanism, it controls the abundance of correctly processed, functional mRNAs. NMD-sensitive mRNAs contain more exon junctions in the 3΄UTR compared to NMD-insensitive transcripts, and, in contrast to previous studies (61–63), long 3΄UTRs did not emerge as a hallmark for NMD sensitivity in our analysis: If we exclude mRNAs with exon-exon junctions in the 3΄UTR, the mean length of the 3΄UTR is similar between NMD-sensitive and insensitive mRNAs, but these remaining NMD-sensitive mRNAs have more uORFs in the 5΄UTR compared to isoforms that are not NMD sensitive.

A major caveat of prior analyses of alternatively spliced transcript isoforms of human genes is that they cannot be unambiguously assembled or accurately quantified from short reads, especially when their abundance is low, as is generally the case with NMD-sensitive transcripts (29,64,65). Full-length transcript sequencing provides a more accurate reference transcriptome that can be used to address this problem, and it has been reported that a vast number of additional isoforms can be detected using this technology (33,36,37,66,67). Recently, a long-read sequencing dataset was obtained from plants depleted of the conserved NMD factor SMG1 (68), although no analysis of this data is currently available. In lung cancer cells, Nanopore sequencing was also used to evaluate the impact of knocking down UPF1 on specific mRNA targets (22). This analysis revealed a significant increase of NMD isoforms and reported novel splicing events for specific, disease-related RNA isoforms. In our study, we use long-read sequencing to perform a more general characterization of NMD-sensitive mRNAs.

Several observations highlight the advantages of our approach in identifying new NMD targets. Approximately 13% of the short-read sequences obtained upon KD of NMD factors mapped exclusively to transcripts that were newly identified by the Nanopore sequencing. Interestingly, splicing level changes are also observed in genes with no significant changes in their overall expression levels. Thus, the number of NMD-sensitive mRNAs identified here is further increased by those cases that were previously missed in analyses that relied on gene-level changes in expression (38,69,70).

The mRNAs that were stabilized by NMD inhibition derived mostly from exon-skipping events. This is in agreement with reports that animals have unusually high rates of alternative exons (71), likely with distinct evolutionary implications (72). For most genes with differences at the expression levels between control and inhibited NMD conditions, only specific isoforms are enriched upon NMD depletion. It is important to elucidate the NMD-sensitive isoforms of a gene, because alternative isoforms may encode distinct proteins that often exert different functions and phenotypical influence, as observed, for example, in the correlation between aberrant transcripts as potential templates for producing neo-antigens in lung cancer cell lines (22).

Some of the advantages of long-read sequencing to detect NMD-sensitive isoforms are illustrated in the case of UTP11 (Fig. 3B). Nanopore sequencing revealed two different NMD-sensitive isoforms, each of which contains a different poised exon. Not only are these exons novel and only detectable upon transcript stabilization by NMD inhibition, but the long-read data shows that the two exons correspond to independent NMD-sensitive isoforms. This approach can be routinely used to analyze the features of NMD targets of interest.

In mammals, the best-characterized NMD-triggering feature is the presence of an exon junction complex (EJC) located more than 30 nucleotides downstream of the termination codon. EJCs are usually deposited 20-24 nucleotides upstream of the splice junctions (73) and stripped off by elongating ribosomes during translation (55,74–76). EJCs that are deposited further downstream of the termination codon are therefore not removed by ribosomes and trigger NMD (77,78). In agreement with this model, our analysis revealed a clear enrichment of mRNAs with a termination codon in the last exon for non-NMD substrates compared to NMD-sensitive mRNAs. As an alternative approach to assess the transcriptome of spliced mRNAs prior to translation, Kovalak and colleagues captured newly synthesized mRNAs by tandem immunoprecipitation of epitope-tagged and untagged EJC components, a technique known as RNA/protein immunoprecipitation in tandem (RIPiT) (79). This analysis revealed that EJC-bound RNAs are highly enriched for spliced transcripts that are subject to rapid clearance by translation-dependent decay (80). Our analysis points to the same direction, because after inhibiting NMD, we enriched for mRNAs with exon junctions in the 3΄UTR that likely still contain EJCs.

However, there are also mRNAs with exon junctions in the 3΄UTR that evade NMD. This agrees with previous reports according to which genetic variations predicted to induce NMD had, in fact, no effect on mRNA levels (81–83). This could be attributed either to the lack of EJCs on certain exon junctions (84) or, most likely, to the fact that additional mRNA or mRNP features are contributing to NMD activation on certain mRNAs.

Several additional features have been reported to stimulate NMD *in vivo*, including long 3΄UTRs (49,85), uORFs, and long exons (7). Our analysis showed that, once exon junctions in the 3΄UTR are controlled for, the length of 3’UTRs is not significantly different between NMD-sensitive and insensitive mRNAs, even though there is ample evidence for reporter mRNAs that become NMD-sensitive by increasing the length of the 3΄UTR (61,62,86–88). It cannot be excluded that it might not be the merely the physical length of the 3΄UTR but rather the mRNP composition that renders these mRNAs sensitive to NMD (2,4). Notably, in our previous study that was based on short-read sequencing reads, we observed that NMD-targeted genes have overall a longer 3΄UTR but with a limited statistical significance (38). Additionally, the exact coding potential of 3΄UTRs must be carefully considered, especially in the light of evidence that alternative splicing can extend the ORFs into the 3΄UTRs (89).

In our dataset, we found the presence of an uORF to be an orthogonal NMD-inducing feature, characteristic of transcripts that do not contain exon-exon junctions in their 3’UTRs. We hypothesize that in cases where the substantial translation of uORFs leads to termination events that are not accompanied by re-initiation, the corresponding mRNAs may be indeed sensed by the NMD mechanism. However, was shown recently that mammalian translation initiation factors eIF4G and eIF3 remain bound to elongating ribosomes and facilitate rescanning after translation of the uORF (90). Additionally, it should be taken into consideration that many computationally predicted uORFs are in fact not translated and therefore do not stimulate NMD (52).

Stochasticity is an innate characteristic at all steps of gene expression that often leads to the production of inert or deleterious RNA molecules (91). Splicing, in particular, is a well-known source of such variation: sequence motifs that resemble those of splice sites are found in intronic regions, leading to aberrant exon inclusions (58,92). To address whether NMD rids the cells of aberrantly spliced mRNAs and thereby fulfills a quality control function, we assessed the 5΄ and 3΄ splice-site scores and conservation rate of NMD-triggering exons compared to exons that do not lead to NMD. This analysis showed that NMD-sensitive mRNAs derive from both correctly and aberrantly spliced mRNAs, with the bulk of events involving weaker and less conserved splice sites. These results favor the view that the primary role of NMD is to degrade aberrantly spliced mRNAs arising from splicing errors. If selection for decreasing the load of aberrant transcripts operates, as would be expected, these events are probably underrepresented even in our currently most comprehensive reference transcriptome (7). On the other hand, this innate variation of gene expression products provides the cells with the flexibility to adapt to environmental cues and can be crucial in developmental processes that are known to be influenced by the NMD activity (91). For this reason, it would be informative to perform a similar analysis in cells of different tissues and developmental stages.

A potential caveat of our experiments is that we cannot distinguish with certainty whether the enriched mRNAs are direct NMD targets or whether they result from altered pre-mRNA splicing due to changes in the concentration of splicing factors, which are often regulated by NMD (20,52). However, given the behavior of the transcripts as well as their features, known to be indicative of NMD, we believe that the majority of the upregulated mRNAs are direct NMD targets.

## Conclusions

Long-read sequencing proved to be highly valuable for identifying NMD-sensitive mRNAs under conditions of reduced NMD activity. The data presented here portray the diversity and multifaceted features of NMD-sensitive mRNAs and provide important information about the splicing origin, the features of these mRNAs as well as their biological role. We expect that this work will serve as an important source to investigate NMD targets at the transcript level and stimulate the usage of long-read sequencing to study the features of RNA degradation substrates in the future. Understanding the mRNA features that lead to NMD is important for understanding the impact of disease-related genetic mutations on the transcriptome and must be taken into consideration in the designing therapeutic schemes, disease models, and genome editing experiments (93,94). Given the current speed of increasing sequencing depth that Nanopore sequencing technology can achieve, we envision that it might soon become feasible to accurately quantify transcript levels directly from long-read sequencing.

## Methods

### Cell culture and knockdown experiments

HeLa cells were cultured in Dulbecco’s modified Eagles medium (DMEM) supplemented with 10% FBS and penicillin/streptomycin and maintained at a temperature of 37°C in a humidified incubator with 5% CO2. Knockdown experiments were performed in HeLa cells as described in (38). Briefly, 6 × 10^5^ HeLa cells were seeded into Τ25 flasks, and 24 h later and the cells were transiently transfected using Dogtor (OZ Biosciences). For single-factor knockdowns, 1 μg of pSUPERpuro plasmids expressing shRNAs against UPF1, SMG6, SMG7, or control plasmids were transfected. For double knockdown experiments, 1 μg of each pSUPERpuro plasmid was added. The cells were split into a T25-cm^2^ cell culture flask and selected with puromycin at a concentration of 1.5 μg/μL. Twenty-four hours prior to harvesting, the cells were washed with PBS and the puromycin-containing medium was exchanged with normal DMEM–FCS medium. Cells were harvested 4 days after transfection. The shRNA target sequences for UPF1 and SMG6 were given in (95) and for SMG7 in (96). The rescue experiments that were used to produce short read sequencing data are described in (38).

### Total RNA isolation and western blot analysis

Total RNA was extracted using the GenElute Mammalian Total RNA Miniprep Kit (Sigma-Aldrich) followed by polyA+ mRNA isolation using Dynabeads Oligo(dT)25 (Thermo Fisher) according to the manufacturer’s guidelines. Cell harvesting for protein samples (derived from the same sample as RNA preparation) was done as previously described (97). Briefly, 2 × 10^5^ cell equivalents were analyzed on a 10% PAGE, and detection was performed using Anti-RENT1 (UPF1) (Bethyl, A300–038A), anti-EST1 (SMG6) (Abcam, ab87539), Anti-SMG7 (Bethyl, A302–170A), and anti-Tyr-Tubulin (Sigma T9028, 1:5000) antibodies.

### Reverse transcription and PCR

Reverse transcription for PCR analysis was performed using 1 μg total RNA and 300 ng random hexamers using Affinity script multiple temperature reverse transcriptase (Agilent). For UTP11, PCR reactions were performed using isoform-specific primer pairs for the two NMD-sensitive isoforms (pair 1: EK158 TGGTGCTTAAAGAGATTGAATAGTATATCCA and EK159 AACGTTTTCGACGACTCTGAAATT, pair 2: EK159 AACGTTTTCGACGACTCTGAAATT and EK160 TTATGGCTCACTGCA GTCTCCA) and a pair of primers that detects the protein-coding isoform of the UTP11 gene (EK161 ATGGCGGCGGCTTTTC and EK159 AACGTTTTCGACGACTCTGAAATT). BAG1 was amplified by PCR using a protein-coding-specific pair of primers (EK141 ACTCATATTTAAGGGAAAATCTCTG / EK38 TTGGGCAGAAAACCCTGCTG) and a pair of primers of the NMD-specific isoform (EK139 CATATTTAAGGTTCTTCAACAGATA / EK140 TGTTTCCATTTCCTTCAGAGA). PCR reactions were performed using CloneAmp™ HiFi PCR Premix (Takara) for 35 cycles with an annealing temperature of 52 °C according to the manufacturer’s suggestions. The total product was run in 1% agarose gels in TAE buffer, stained with ethidium bromide and visualized under UV light.

### Library preparation and Oxford Nanopore sequencing

The Oxford Nanopore Technology SQK-DCS108 kit was used for PCR-free direct cDNA sequencing. For each DCS108 library (CTR, UPF1, SMG6, SMG7, CTR, dKD), 500 ng of polyA+RNA was used as input, following the instructions of the ONT protocol. The libraries were sequenced on a GridION using R9.4 flowcells and with the MinKNOW instrument software, we recorded the Nanopore current and generated fast5 files. Basecalling of the fast5 files was performed using GUPPY version (1.4.3-1 from Oxford Nanopore Technologies).

### Transcriptome generation from Oxford Nanopore reads

The human reference genome and annotation were obtained from ENSEMBL (40) (version 90/ hg38). Oxford nanopore reads that were obtained from the GUPPY base caller in .fastq format were mapped to the reference genome using minimap2 (39) (version 2.17) with the option ‘-ax splice’. Alignments were sorted and indexed with samtools (98). Stringtie (99) (version 2.0) was used to construct a transcriptome for each of the individual samples with the options ‘-G <reference annotation> -L’. The sample-specific annotation files were merged with Stringtie with the options ‘--merge -G <reference annotation>’. CDS prediction was performed with a custom python script that relies on the following libraries (HTSeq (100) (version 0.10.0), BioPython(101) (version 1.71) and PyFasta (102) (version 0.5.2)). The tool selects one start codon for each gene from the APPRIS (103) database of high-confidence transcript isoforms and then looks for a downstream in-frame stop codon in the transcripts annotated for the given gene.

### Processing of Illumina reads

Reads were trimmed with the trimFastq.py script of rMATS (43) to obtain reads of 75 nts, as required for the splicing analysis. A genome index was built with STAR (version 2.7.1a) (42) using the annotation derived from long-read sequencing, and the reads were mapped again with STAR to the reference genome, with the following options ‘--alignEndsType EndToEnd --outSAMstrandField intronMotif --alignIntronMax 1’. Differential Splicing Analysis at the event level was performed with rMATS(43) (version 4.0.2) with default options. For each event type (skipped exons, retained introns, mutually exclusive exons, alternative 5΄/3΄ splice sites) rMATS gives a percent spliced in (*PSI*) score, indicating the proportion of transcripts that provide positive evidence for the event relative to all transcripts whose pre-mRNAs cover the genomic region of the event. *PSI* scores are used to calculate also ‘Δ*PSI*’ values, comparing two distinct conditions. Salmon (44) (version 0.14.1) was used to estimate the expression levels of genes and transcripts, while edgeR (45) (version 3.22.3) was used for differential expression analysis at the gene and transcript level.

### Selection of NMD-responding events

To identify NMD-sensitive events, we used the short-read-based *PSI* values obtained from control, dKD and rescue samples, and the associated Δ*PSI* values generated by rMATS for comparisons of double knockdown with control and rescue with double knockdown samples. Events with FDR < 0.05 in both comparisons, with the additional restriction of at least 10% change (in absolute value) in the dKD vs. control comparison and a change in the opposite direction in the rescue vs. dKD comparison. For each category of events, we also distinguished two subclasses, depending on whether the ‘inclusion’ or ‘exclusion’ form of the event is enhanced when NMD is repressed.

### Selection of transcripts related to individual events for gene and isoform expression analysis

To evaluate the effect of NMD on transcript and gene expression, we identified all transcripts related to significant events, in particular alternative exons, based on the corresponding pre-mRNA covering the genomic locus of the event. For events where the inclusion was associated with NMD, we selected the transcripts that contained the exon while for the events where the exclusion was associated with NMD, we selected the transcripts that did not include the exon.

### Selection of NMD-sensitive and insensitive isoforms for individual genes for feature analysis

We started again from the significant events and identified all transcripts related to these events, as described above. We filtered out those transcripts without annotated coding regions and then separated the remaining transcripts into potentially NMD-sensitive or insensitive, depending on which form of the event was enhanced by the repression of NMD. That is, in the case of skipping exons, if the *PSI* was higher in the double knockdown compared to control samples, we considered the transcript including the exon to be NMD-sensitive and those that did not include the exon to be NMD-insensitive. Finally, to obtain a non-redundant set of transcripts, we further selected from each subset the transcript with the highest average abundance in control and double knockdown samples. These were used to determine the number of exons, 3’ UTR length, etc. For the uORF analysis, we identified transcripts that contain an uORF defined by a start (AUG) and stop (UAA, UGA, UAG) codons, with a length of at least 15 nts (including the stop codon) and were located entirely upstream of the main annotated ORF. To focus on uORF that are indeed translated, we intersected the computationally predicted uORFs with recently published RiboSeq data (57) and retained only those with experimental evidence for translation. This analysis was performed with zgtf of the ZARP pipeline (104).

### Features of NMD-inducing skipping exons

We compared various features of NMD-inducing skipping exons and randomly chosen internal exons. The ‘background’ set of exons was chosen dependent on the feature. To compare exon length, we used all internal exons in transcripts with an annotated coding region. To compare splice signals and evolutionary conservation, we chose the same number of internal exons from any transcripts with an annotated coding region as we had NMD-inducing transcripts. We further imposed the restriction to have only one internal exon per gene.

To calculate the strength of splice sites, we extracted 20 bases in the intron and 3 bases in the exon around a 3’splice site, and 3 bases in the exon and 6 bases in the intron around a 5’splice site. We then submitted these sequences to MaxEntScan (59) to obtain scores reflecting the strength of these splice sites, which we compared between NMD-inducing and background exons. Similarly, we extracted PhyloP30 scores (60) for the above-mentioned regions around 5’and 3’splice sites. The average scores per position in each of these windows were calculated and we compared the distributions of these scores between NMD-inducing and background exons.

## Supporting information

Table S1 Pairs of NMD-sensitve/insensitive isoforms

## Abbreviations

NMD: Nonsense-mediated mRNA decay
AS-NMD: Alternative splicing linked to NMD
EJC: Exon junction complex
KD: Knockdown
dKD: Double knockdown
TC: Termination codon
PTC: Premature termination codon
PSI: Percent spliced in
ORF: Open reading frame
uORF: Upstream open reading frame

## Declarations

## Acknowledgments

We thank Alban Ramette and his group from the University of Bern for technical help in the first Nanopore sequencing experiments.

## Availability of data and materials

The Illumina sequencing datasets analyzed in this study were downloaded from GEO (GSE86148) (38). RNA-seq data have been deposited in the ArrayExpress database at EMBL-EBI (www.ebi.ac.uk/arrayexpress) under accession number E-MTAB-10452.

## Authors’ contributions

EK and OM conceived the project. EK performed the experiments, constructed the cDNA-Seq libraries and wrote the manuscript. FG and MZ performed the bioinformatics analysis. All authors contributed to the data analysis and interpretation, and to the final editing of the manuscript.

## Ethics approval and consent to participate

Not applicable.

## Consent for publication

Not applicable

## Supplementary Figures

**Supplementary Figure 1.**
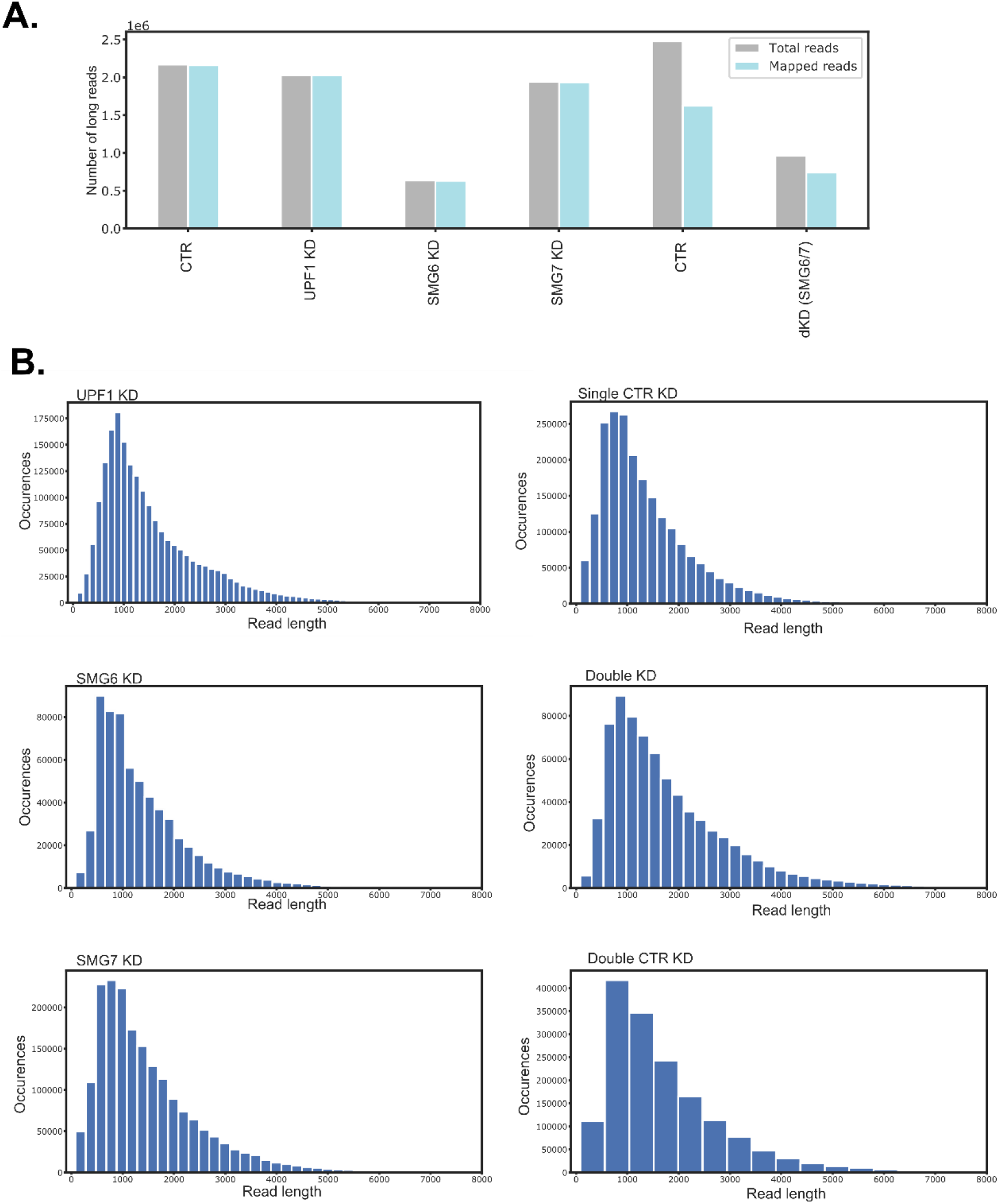
**(A)** Number of totally sequenced (grey) and mapped Nanopore reads (light blue) in the different cDNA sequencing experiments of reverse-transcribed total RNA from cells with the indicated knockdowns. **(B)** Read length distribution of mapped reads from the long-read cDNA sequencing experiments indicated in (A).

**Supplementary Figure 2.**
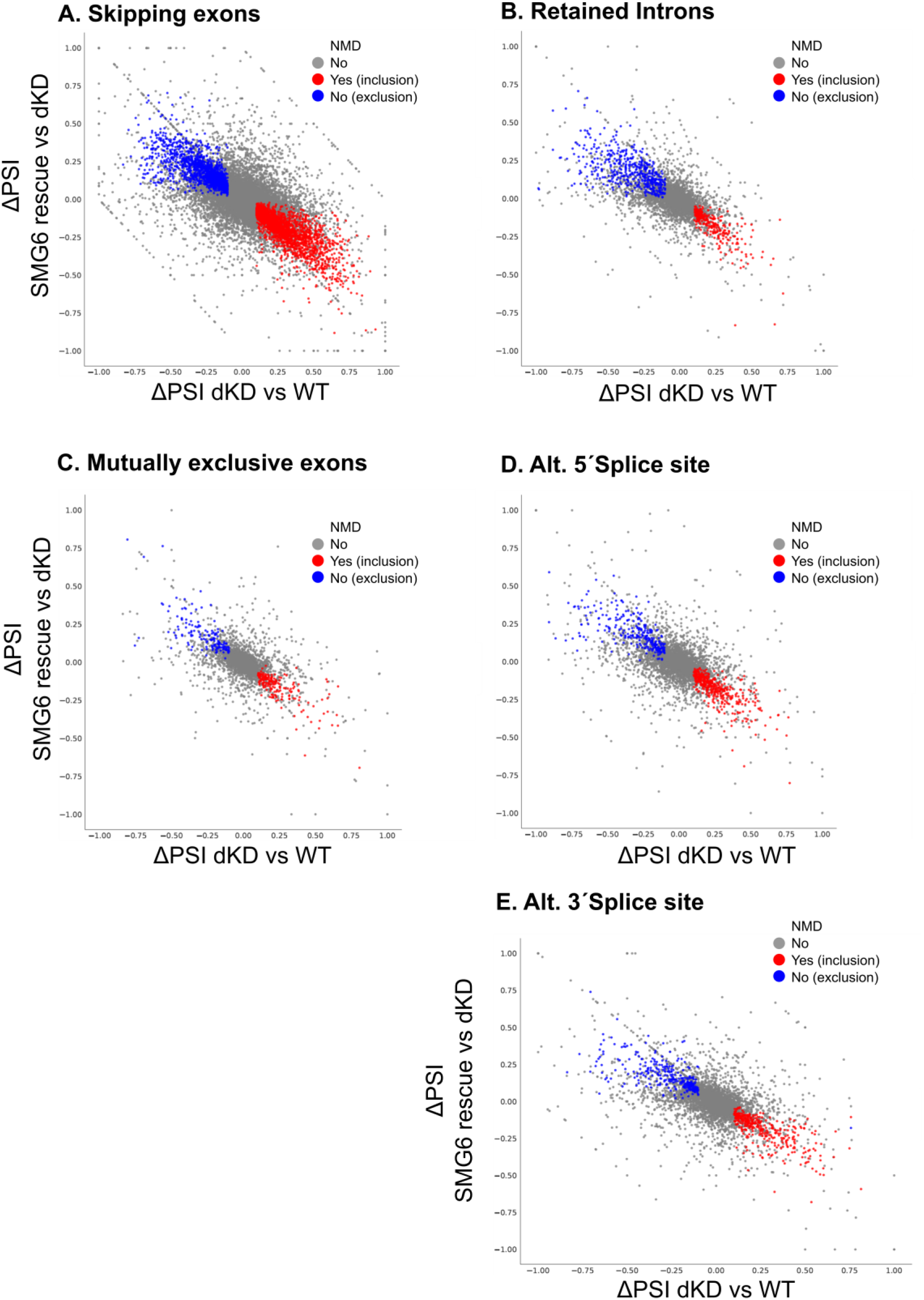
(A-E) Scatter plots comparing the change in “percent spliced in” (deltaPSI) of exons in CTR versus dKD (*x*-axis) and SMG6 rescue versus dKD (*y*-axis). Coloured in red are significant exon inclusion and in blue significant exon exclusion events. Non-significant events are depicted in grey.

